# Mind the gap: A systematic review and meta-analysis of how social memory is studied

**DOI:** 10.1101/2023.12.20.572606

**Authors:** Meghan Cum, Jocelyn Santiago Pérez, Erika Wangia, Naeliz Lopez, Elizabeth S. Wright, Ryo L. Iwata, Albert Li, Amelia R. Chambers, Nancy Padilla-Coreano

## Abstract

Social recognition is crucial for survival in social species, and necessary for group living, selective reproduction, pair bonding, and dominance hierarchies. Mice and rats are the most commonly used animal models in social memory research, however current paradigms do not account for the complex social dynamics they exhibit in the wild. To assess the range of social memories being studied, we conducted a systematic analysis of neuroscience articles testing the social memory of mice and rats published within the past two decades and analyzed their methods. Our results show that despite these rodent’s rich social memory capabilities, the majority of social recognition papers explore short-term memories and short-term familiarity levels with minimal exposure between subject and familiar stimuli – a narrow type of social memory. We have identified several key areas currently understudied or underrepresented: kin relationships, mates, social ranks, sex variabilities, and the effects of aging. Additionally, reporting on social stimulus variables such as housing history, strain, and age, is limited, which may impede reproducibility. Overall, our data highlight large gaps in the diversity of social memories studied and the effects social variables have on social memory mechanisms.

## Introduction

Social memory is critical for individuals to adaptively navigate a social world. Social memory in humans has been a topic of study since the 1950s^1,2^, and in rodents, experimental exploration spans back to the 1970s^3,4^. Rodents naturally demonstrate the ability to use social memory to navigate their surroundings. Wild rodents outside of the laboratory form complex social groups that display the ability to discriminate between dozens of individuals. For example, wild house mice interact with dozens of individuals during a month’s time and form social structures to control territory and selectively mate with other high-ranking individuals^5^. Similarly, rats outside the laboratory form complex and stable social structures that require distinguishing between large number of individuals for long periods of time^6,7^. Laboratory and wild mice both demonstrate social recognition of kin^8^. Furthermore, in captivity, both wild mice^9,10^ and laboratory mice^11,12^ form social hierarchies. In a large vivarium of 30 individuals, mice formed two distinct dominance subnetworks that were stable across weeks, showcasing their ability to show social preferences and distinguish among many familiar individuals^13^. Laboratory mice and rats make up the bulk of all neuroscience research and thus the bulk of social memory research^14^. Individuals of both species live in groups and show long-term relationships that matter for their behavior in their natural environment, thus making these long-term relationships ethologically relevant (Table 1). These rich long-term relationships suggest that the identity of individuals and types of relationships is encoded in the brain. Understanding how the brain encodes identity and distinct types of relationships is important for our fundamental understanding of social cognition. In addition, studying diverse and long-term social memories has strong clinical implications given that rodents are used as models to generate treatments for social and memory deficits associated with neurological and neuropsychiatric disorders^15,16^.

**Table 1:** Definitions relevant to this study.

| Terms | Definitions |
| --- | --- |
| **Ethologically relevant relationship** | A social relationship that plays specific roles based on the nature of a species. Short term social memories that could be ethologically relevant are recognizing intruders or predators. Social memories of kin, mate, or members of a social hierarchy are examples of long term ethologically relevant memories. |
| **Familiarity level** | Amount of exposure time the subject has had with a particular individual or social stimulus. |
| **Long-term familiarity** | Exposure between subject and social stimulus is more than 1 day. |
| **Short-term familiarity** | Exposure between subject and social stimulus is less than 1 day. |
| **Long-term memory** | Social memories that have at least 24 hours since encoding began. |
| **Short-term memory** | Social memories that have less than 24 hours since encoding began. Of note some studies use even shorter windows of less than 6 hours to define short term memory |
| **Social stimuli** | Conspecifics used in social interactions with the subject. They can be novel or familiar animals depending on the experiment. |

Within the laboratory, a few paradigms are predominantly used to test social memory. In each paradigm there is a subject, the animal whose behavior is being measured, and a social stimulus, the animal being presented to the subject. Rats and mice have been shown to have both sociability preferences, (preferring social interaction to no social interaction), and social novelty preferences (preferring interaction with a novel vs. familiar animal)^17,18^. Behavior such as less investigation time across repeated exposure to the same social stimulus or more investigation or a novel versus a familiar social stimulus suggests social recognition of the subject to the stimulus. The paradigms that have mainly been used, with various modifications, include: the habituation-dishabituation paradigm, the five-trial social memory test (**Figure 1a**), and the three-chamber social assay (**Figure 1b**), all of which have been described in detail elsewhere^18–20^. Time exploring the social stimuli decreases as familiarization increases in the habituation-dishabituation paradigm and time in the chamber with the novel stimulus is higher in the three-chamber assay. More recently, given the limitations of the three-chamber assay^21^, there have been smaller chambers used to present a familiar and novel stimulus reducing the need for spatial navigation^22,22–24^ (**Figure 1c**). In addition, a handful of studies have created operant based tasks (**Figure 1d**) to study social memories using two familiar individuals^17,25,26^, as well as variations in the social recognition paradigm in order to present more social stimuli (**Figure 1e**) and measure behavior in a more data-driven way^27^.

**Figure 1.**
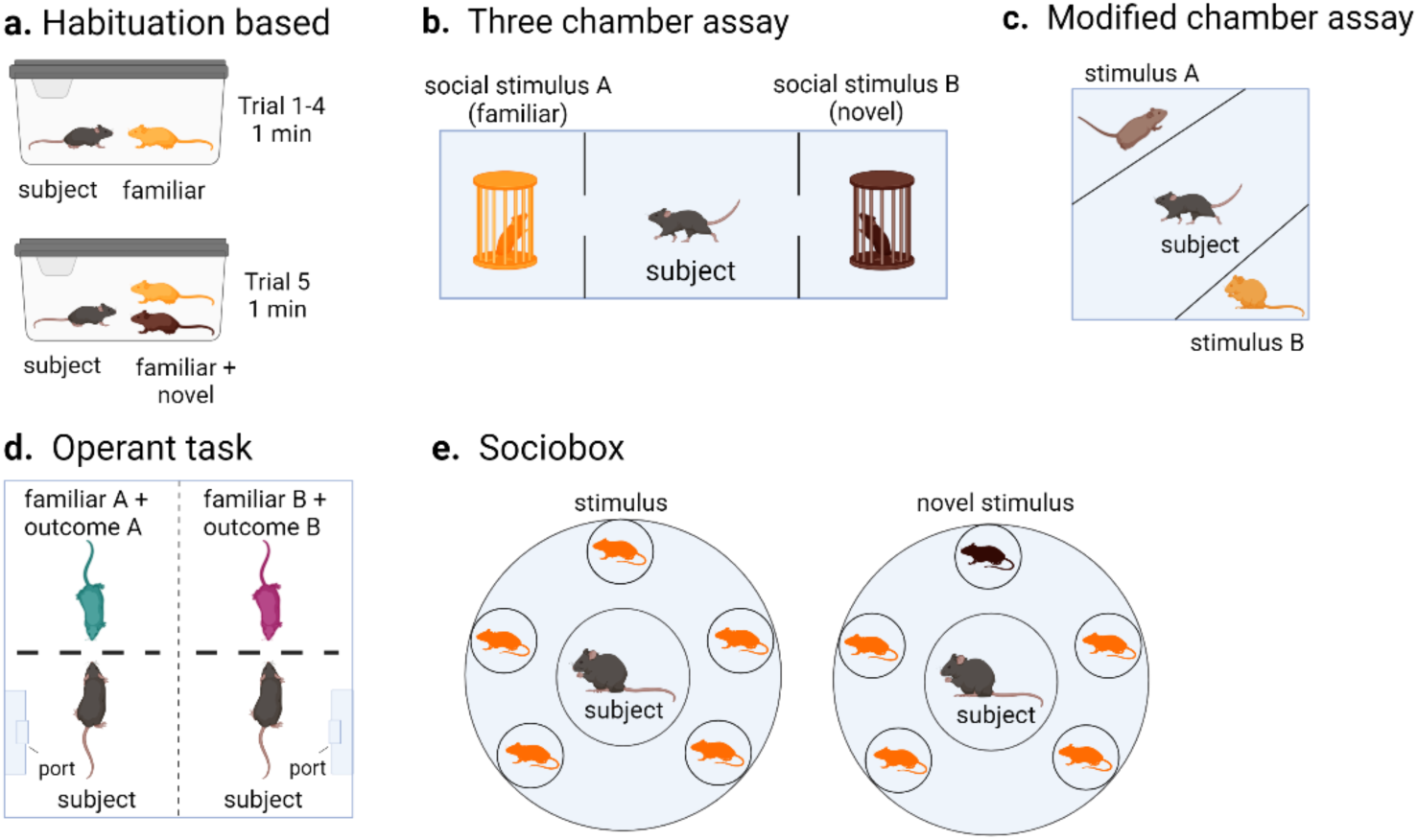
Social memory paradigms. **a.** Schematic showing standard trial-based social memory assays. **b.** Schematic of classic 3-chamber social memory assay. **c.** Modified chamber assay apparatus **d.** Operant task where subject learns to associate familiar social stimuli with a non-social outcome; social discrimination is judged based on subject’s learned behavioral output. **e.** Sociobox apparatus designed to test social memory against multiple present familiar social stimuli. Left shows phase one for familiarization and right is social recognition phase.

Social interactions between individuals vary depending on social history and other biological factors (Table 2). For example, male mice and rats have been shown to have higher levels of baseline social stimulus investigation as compared to females^28,29^, but female rats have been shown to retain social recognition for longer periods of time^30^. Rodents can exhibit social recognition up to 3 -7 days later after a two-minute exposure to a social stimulus^31–33^ while, paternal male mice can remember their offspring 5-6 weeks after separation ^34^. Considering the many social relationships that mice and rats have and the wide use of rodent models, we quantified the types of memories studied and how they are being tested in the neuroscience field. To help shape future social memory research, we systematically quantified gaps that exist in the current literature and outline factors and suggestions to fill these gaps in future social memory studies.

**Table 2:**
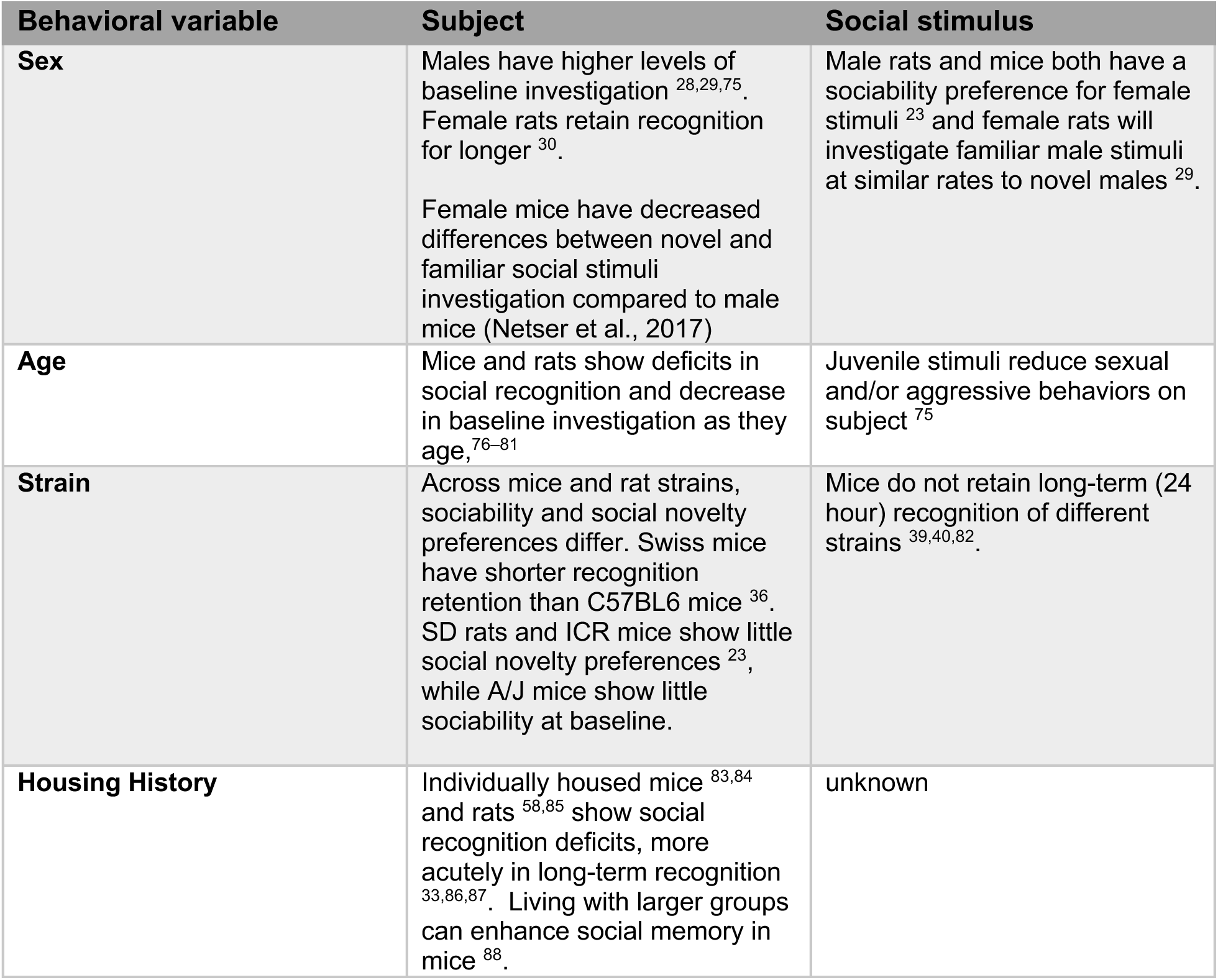
Evidence of social investigation metrics changing with social and biological variables.

| Behavioral variable | Subject | Social stimulus |
| --- | --- | --- |
| **Sex** | Males have higher levels of baseline investigation <sup>28,29,75</sup>. Female rats retain recognition for longer <sup>30</sup>.<br>Female mice have decreased differences between novel and familiar social stimuli investigation compared to male mice (Netser et al., 2017) | Male rats and mice both have a sociability preference for female stimuli <sup>23</sup> and female rats will investigate familiar male stimuli at similar rates to novel males <sup>29</sup>. |
| **Age** | Mice and rats show deficits in social recognition and decrease in baseline investigation as they age, <sup>76-81</sup> | Juvenile stimuli reduce sexual and/or aggressive behaviors on subject <sup>75</sup> |
| **Strain** | Across mice and rat strains, sociability and social novelty preferences differ. Swiss mice have shorter recognition retention than C57BL6 mice <sup>36</sup>. SD rats and ICR mice show little social novelty preferences <sup>23</sup>, while A/J mice show little sociability at baseline. | Mice do not retain long-term (24 hour) recognition of different strains <sup>39,40,82</sup>. |
| **Housing History** | Individually housed mice <sup>83,84</sup> and rats <sup>58,85</sup> show social recognition deficits, more acutely in long-term recognition <sup>33,86,87</sup>. Living with larger groups can enhance social memory in mice <sup>88</sup>. | unknown |

## Methods

We searched four databases: Web of Science, Embase, PubMed, and PsycINFO, all on January 14, 2022. We focused our search on the abstract text, as such the scope of this review is limited by the terminology used by the primary research authors in the abstract (**Figure 2** - search criteria). Our goal was to review those articles where social memory was at the center of the goal of the study, rather than those where social recognition experiments served as controls. Studies were selected by several parameters: 1) the term “social” was used to describe the type of memory, discrimination or recognition being investigated in the abstract, 2) we excluded purely behavioral studies by requiring the use of the term “neuro” in the abstract. While this limited our search, we believe using these boundaries allowed us to focus on studies pertaining to the neural mechanisms of social memory without further author bias to categorize other memory studies. Each paper was reviewed to confirm whether the article satisfied all criteria to be included in this review (**Figure 2**). After each paper was reviewed for criteria satisfaction, 672 qualified articles remained. Each article was analyzed and quantified for various methodological variables. EndNote Online and Zotero were used as reference managers. Detailed results can be found in the supplemental materials.

**Figure 2.**
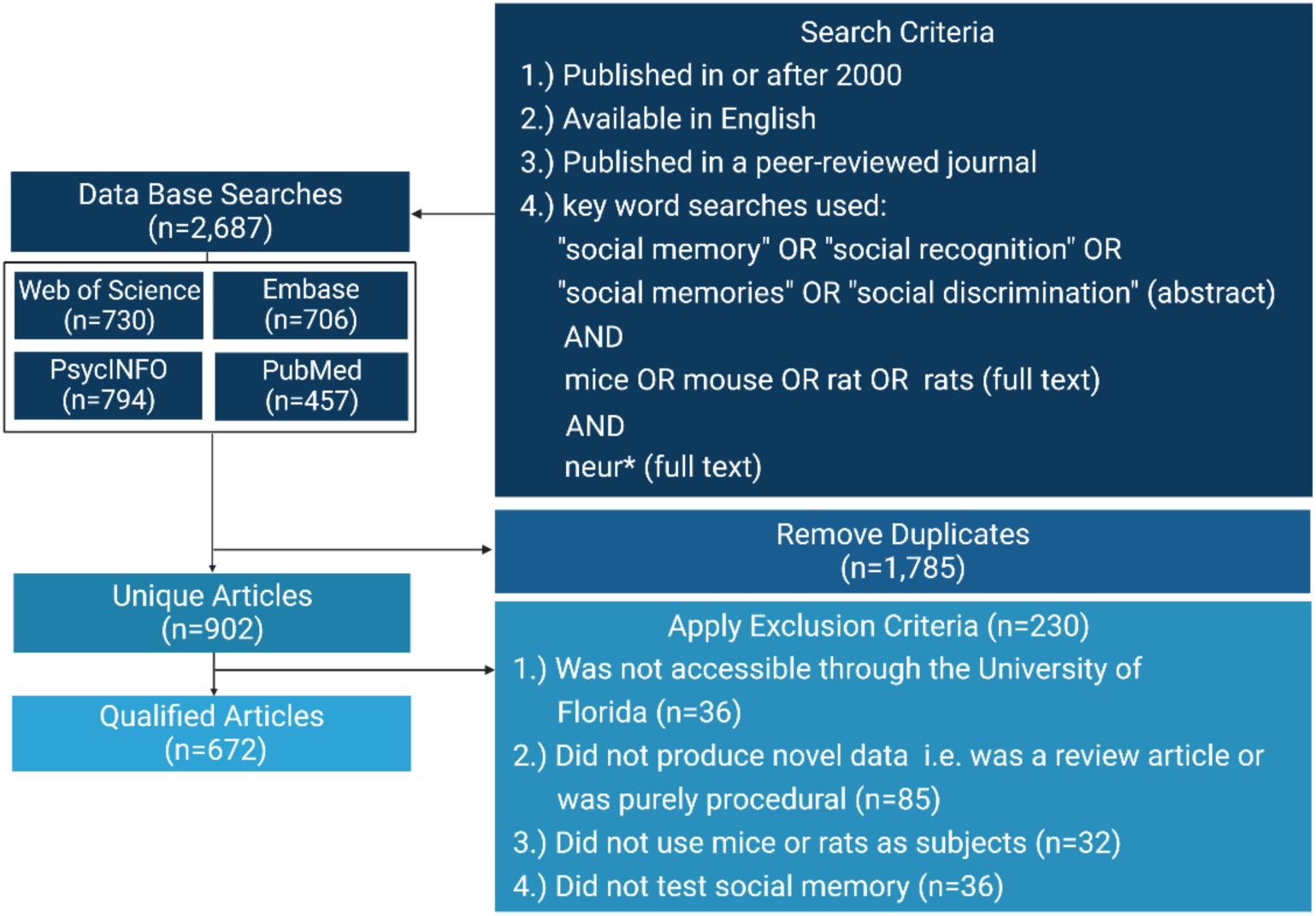
Methodological Flowchart. Databases searched with numbers (*n*) of articles found per site alongside key terms and search criteria, number of duplicates, and list of exclusion criteria with the number (n) of articles removed for each item.

To satisfy our criteria for testing social memory, an article must have included a behavioral experiment consisting of more than one exposure to the same social stimulus or social odor (e.g. bedding or urine from a conspecific). Additionally, the authors must have reported some behavioral metric that indicated social recognition. Any significant behavioral differences due to familiar or novel social stimuli exposure qualified. Sniffing time, exploration time, interaction time etc. were all recorded as social investigation time. If a subject was exposed multiple times to the same social stimulus but no behaviors were measured and compared across exposures to confirm recognition of an individual social stimulus by the subject, the experiment did not satisfy our criteria. Examples of paradigms that did not meet our social memory criteria were chronic social defeat studies (no behavioral metric indicating recognition) and social transmission of food preference (single exposure to a conspecific).

All experimental conditions used in an article were analyzed and a single article can be represented multiple times in a figure if it used different subjects, social stimuli, intertrial intervals, etc. If no ovariectomy procedures were mentioned, female subjects and social stimuli were assumed to be intact. If ovariectomies were mentioned, but no age, female social stimuli were assumed to be adults. Housing conditions were divided into four groups: chronic isolation, acute isolation, group-housed, or not specified. Chronic isolation was defined as being singly housed for longer than a week. Familiarity levels for familiar social stimuli were either categorized by the ethologically relevant relationship to the subject, e.g., littermate, or were defined as the total time of exposure prior to a test trial.

To substantiate that any quantified methodological variable was used significantly more than any other, a two-sample chi-square test was done for the two most used conditions such that the second most used condition was the observed frequency, and the most used condition was the expected frequency. *Other* and *not specified* are not homogenous groups and therefore were not used in any chi-square tests.

## Results

### While social memory research has increased, the subjects remain adult male mice

Social memory, and its underlying neural mechanisms, has been growing as a research topic. The number of yearly published papers more than doubled between 2017 and 2021 (**Figure 3**). We next investigated for each of these articles, what type of subjects, social stimuli and social memories were tested.

**Figure 3.**
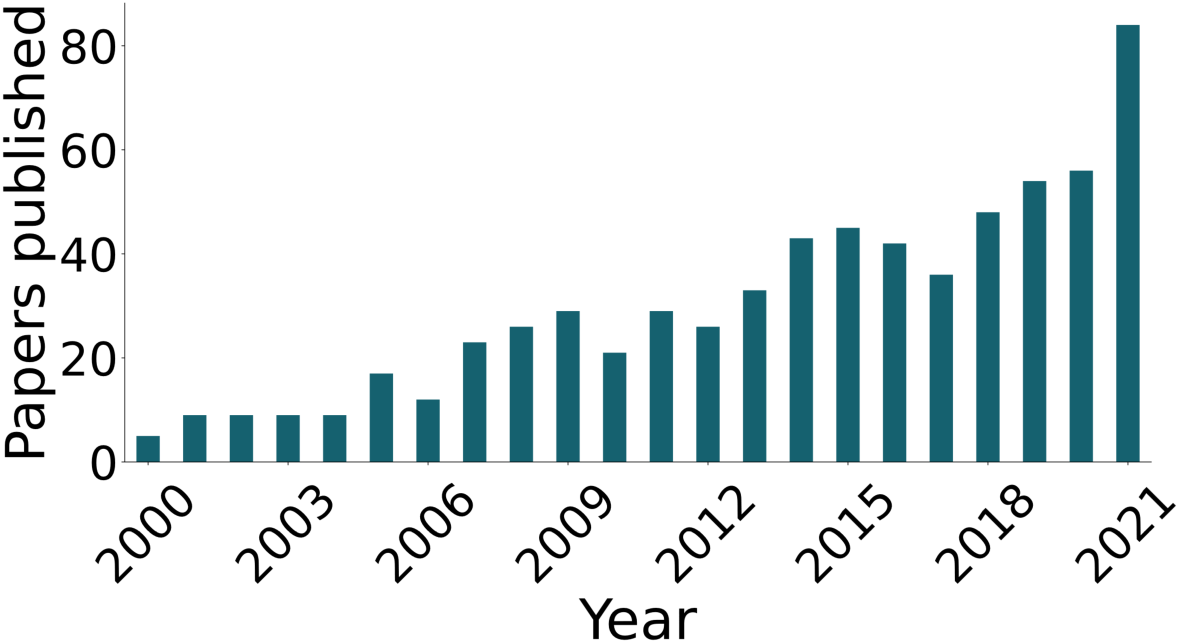
Social Memory Articles Published in the 2000’s. Articles that satisfied our inclusion criteria and were included in this studied plotted by the year they were published.

We defined the subject as the experimental animal whose social memory is being tested by recording their behavior. We found mouse models were more popular than rat models with 60% of articles using mice and 41% using rats, and 1% of articles using both. Given the published work showing social behavior differences across strains of mice and rats ^23,35,36^, we quantified which strains dominate social memory experiments. Mouse research widely used the inbred C57BL6 strain (C57BL6 vs C57BL6 mix, two sample chi-square, X^2^ = 492.97, n = 406, p = 3.22×10^-109^; **Figure 4a**). Amongst the rat studies, two outbred strains, Wistar and Sprague-Dawley, were predominant (**Figure 4b**). When it came to the age of the subjects, very few articles (n = 14 out of 672) used aged subjects (72 weeks and older) for social memory experiments (**Figure 4c**). Although the sex gap in social memory research is decreasing (**Figure 4e**), males still have significantly greater representation than females (female vs male, two sample chi-square, X^2^ = 2048.52, n = 672, p = 0; **Figure 4d**). For housing conditions, group-housing has become the dominant practice since 2006 (**Figure 4g**). Of the 169 articles utilizing chronic or acute isolation, 17% explicitly assessed the effects of social isolation on social memory. For the other studies, many used social isolation to increase social motivation, establish home-cage territory, or to allow surgical recovery. Despite the documented effects of social housing on many behavioral and neurophysiological variables^37^, an average of 18% of articles in the past decade did not report housing conditions for subjects (**Figure 4f**).

**Figure 4.**
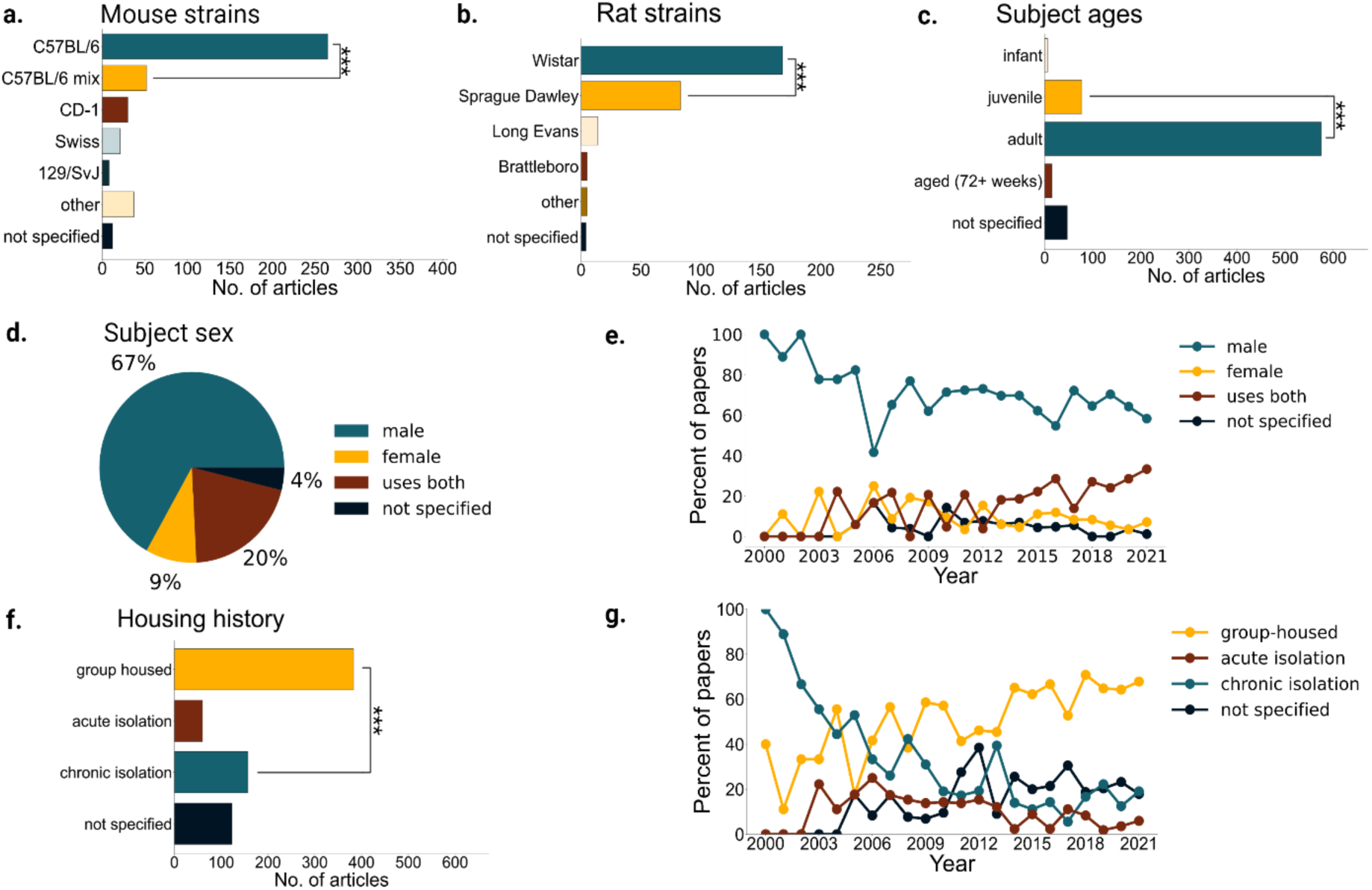
Subject Identity in social memory studies. **a.** Number of articles using different strains of mice (n = 406 from 672 articles). Other strains for mice included: BALB/c, BTBR, A/J, C3H, FVB/NJ, ICR, OF1, NMRI, DBA, CF-1, and wild mice. C57BL/6 mix vs C57BL6 two sample chi-square, X^2^ = 492.96, p = 3.22×10^-109^. **b.** Number of articles using different strains of rats (n = 275 from 672). Other strains for rats included Lewis, SHR, and WAG/Rjj. Sprague Dawley vs Wistar two sample chi-square, X^2^ = 110.53, p = 7.50×10^-26^. Note that the 1% articles that included both rats and mice and are represented in both a and b. **c.** Ages of subjects across all articles reviewed. Infant <3 weeks, juvenile 3-7 weeks, adult >7 & <72 weeks, aged > 72 weeks. Juveniles vs adults two sample chi-square, X^2^ = 3,637.64, p = 0. **d.** Pie chart showing the percentage of all articles (n = 672) from years 2000-2022 that use one, both or unspecified sex of mice or rats. **e.** Percent of articles from years 2000-2021 (n = 665) that use one, both, or unspecified sexes of mice or rats. If no sexes were stated, we marked papers as “sex not specified”. **f.** Number of articles from years 2000-2021 (n = 665) that use group-housing, acute isolation (over a day – less than a week), chronic isolation (over a week), or do not specify the housing history of the subjects. Chronic isolation vs group housed two sample chi-square, X^2^ = 310.09, p = 2.09×10^-69^. **g.** Number of articles from 2000-2022 reporting each type of housing history.

### When described, social stimuli tend to be young males

Social interaction and social memory are modified by the identities or behavioral state of both the subject and the social stimulus^38–40^. Therefore, it is important to consider what types of social stimuli have been used. Despite the potential effects that the identity of social stimulus can have on social interactions (Table 2), we found that most studies lacked information concerning social stimuli characteristics. The strain of social stimuli was often not reported (**Figure 5b, c**) and unless clearly stated in the methods of a given article, we did not assume that the social stimulus was of the same strain as the subject. When information was reported, the most commonly used social stimulus was a male juvenile (**Figure 5a, e**) with males being used considerably more than females (female vs male, two sample chi-square, X^2^ = 333.95, n = 672, p = 1.32×10^-74^; **Figure 4e**). Of the adult female social stimuli, almost half were ovariectomized (**Figure 5f**). We encountered a one article that used ovariectomized juvenile females^41^. Of note, it has been shown that estrogen disruptions in subjects, as is caused by ovariectomies, can cause disruptions in social memory^42–44^. As social interactions are dynamic, where the social stimulus can affect the behavior of the subject^23^, it may be possible that using ovariectomized social stimuli affects the social memory of the subject, but this remains unknown. The most common variable not reported was the housing condition of the social stimulus (**Figure 5d**). Overall, we found that social stimuli characteristics are less reported than those of the subject.

**Figure 5.**
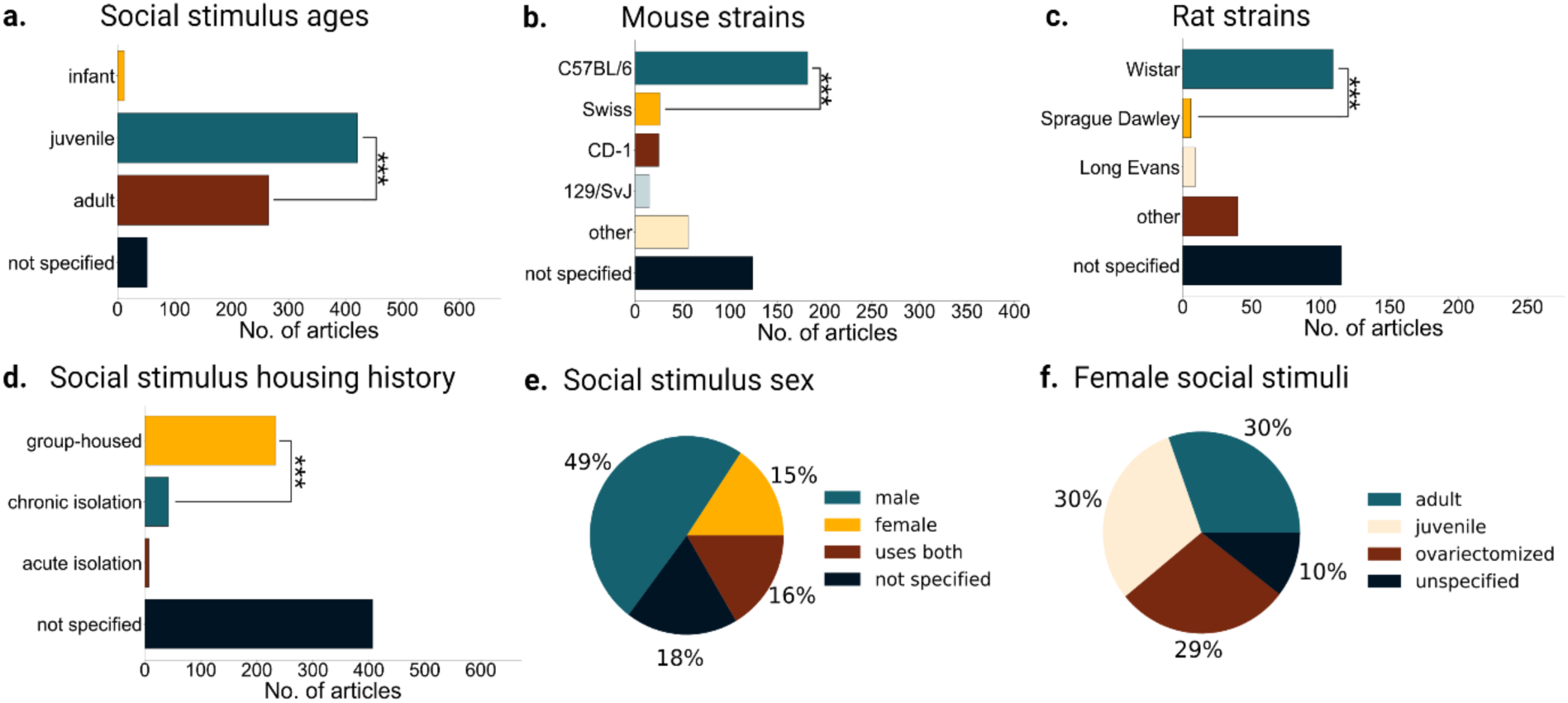
Social stimulus identities in social memory studies. **a.** Age of social stimuli across all articles, adult vs juveniles two sample chi-square, X^2^ = 154.76, n = 672, p = 1.58×10^-35^. **b.** For 406 mouse articles that included social stimuli, mouse strains that were used for social stimuli. Other mouse strains include BALB/c, BTBR, A/J, C3H, FVB/NJ, ICR, OF1, NMRI, DBA, and wild mice. Swiss vs C57BL/6 two sample chi-square, X^2^ = 239.51, p = 5.02×10^-54^. **c.** For 279 rat articles that included social stimuli, rat strains that were used for social stimuli. Other rat strains include Lewis, Lister Hooded, WAG/Rij and F344. Sprague Dawley vs Wistar two sample chi-square, X^2^ = 83.64, p = 5.95×10^-20^. **d.** Number of articles that used group-housing, acute isolation (over a day – less than a week), chronic isolation (over a week), or do not specify the housing history for social stimuli. Chronic isolation vs group housed two sample chi-square, X^2^ = 241.70, n = 672, p = 1.67×10^-54^. **e.** Percent of all articles (n = 672) that use one, both, or unspecified sexes of mice or rats for social stimuli. **f.** Of those articles that used female social stimuli (n = 220), percentage of those that used juvenile, ovariectomized, adult intact or unspecified age females.

### Few studies measure long-term, ethologically relevant relationships

Levels of familiarity between the familiar social stimulus and subject being tested were typically five minutes or less (6-120 minutes vs 5 minutes or less, two sample chi-square, X^2^ = 230.68, n = 672, p = 4.25×10^-52^; **Figure 6a-b**) with most subjects spending at most 2 hours with the social stimulus prior to being tested for social recognition. Of the few articles (n = 25 out of 672) that tested longer-term relationships for social recognition, most studied social recognition of littermates (**Figure 6c).** For parent offspring relationships, 3 studies tested an infant’s ability to recognize the odor or presence of their mother vs. a novel dam^45–47^. Yang et al. tested the offspring’s recognition of its mother in adulthood as well. These studies demonstrate that the capacity for social memory develops early in life and recruits the hippocampal CA2, demonstrating that the role of CA2 in social memory starts early in development and spans both short and long-term social memories. Additionally, by using a foster mother, Laham et al. confirms that offspring’s maternal preferences are not innate genetic preferences but require experience for social recognition. One study demonstrated the ability of a male parent to recognize its offspring after 5-6 weeks of separation^34^. Only one paper tested the memory of a female subject for its mate by measuring pregnancy blocking, a female’s ability to terminate pregnancy in the presence of a novel male^48^ (**Figure 6c**). In this study the authors showed that the memory for the mate is dependent on oxytocin action, demonstrating some potential overlap with social memory mechanisms and pregnancy blocking. Studies that use ethologically relevant long-term relationships enrich our fundamental knowledge of social memory and could facilitate identifying mechanisms that are distinct for short vs long-term familiarity. Despite the importance of social hierarchies and of the effects of social rank in behavior and brain function, most social memory studies do not consider social rank. Given the literature of the increased attention subordinates give towards dominant animals^49^, it is likely that the neural mechanisms of long-term social memory are modulated by social rank to facilitate quick behavioral discrimination of social rank. Of all the articles that used group-housed adult rodents, only one article (**Figure 6c**) studied the interplay between social rank and social memory. Cordero and Sandi found that stress potentiates long-term maintenance, or memory, of a social hierarchy ^50^. Our results highlight the lack of inclusion and consideration in the past literature for long-term ethologically relevant social memory mechanisms.

**Figure 6.**
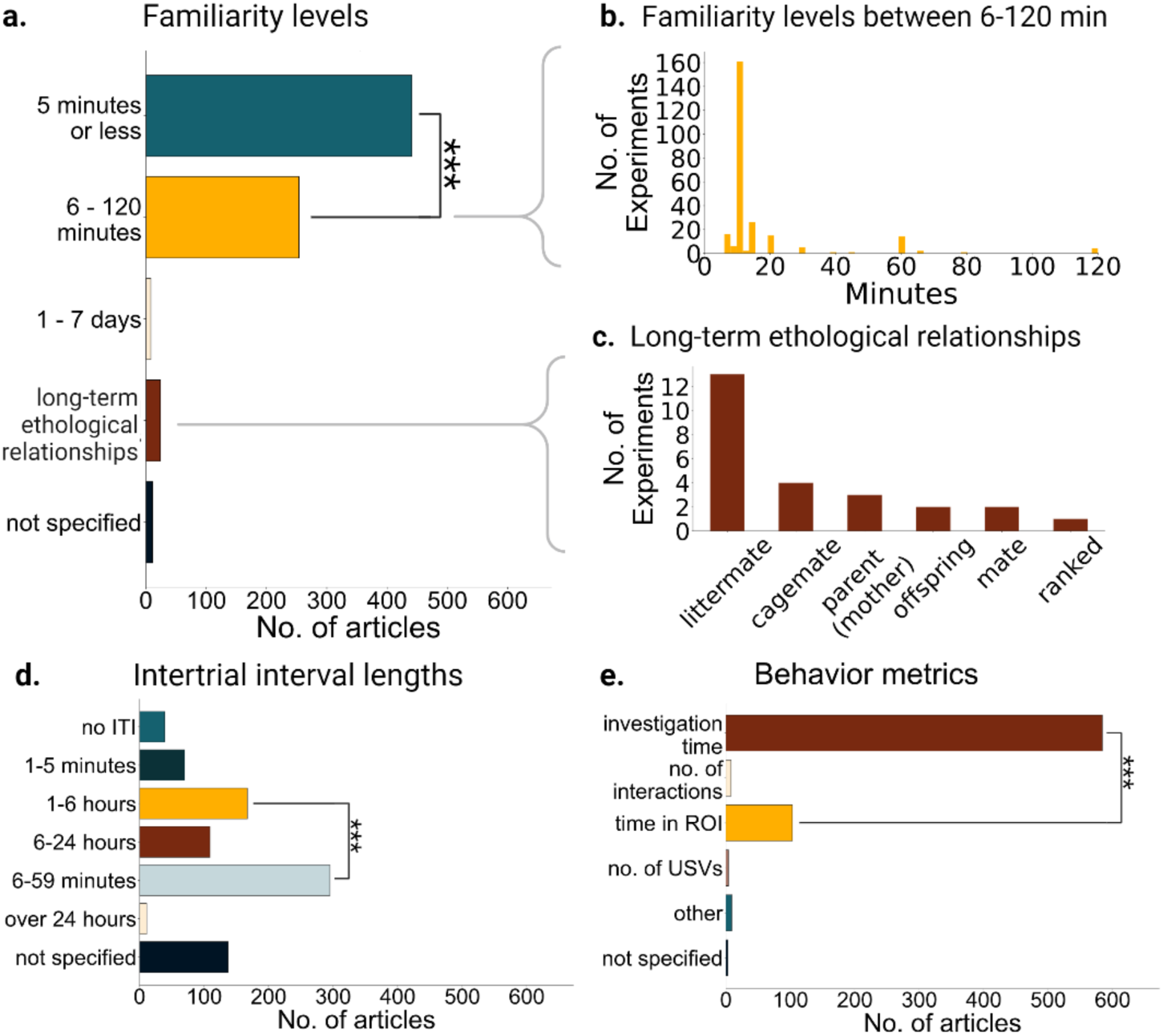
Types of social memories studied and how they were measured. **a.** Number of articles that use various levels of familiarity. 6-120 minutes vs 5 minutes or less two sample chi-square, X^2^ = 230.68, n = 672, p = 4.25×10^-52^. **b.** Histogram showing all familiarity levels used from 6 to 120 minutes. **c.** A breakdown of all the types of ethologically relevant relationships tested in a social memory paradigm in n = 25 articles **d.** Number of articles that use various intertrial interval (ITI) lengths; ITI being the time in between trials, both learning and test trials. 1-6 hours vs 6-59 minutes two sample chi-square, X^2^ = 99.00, n = 672, p = 2.53×10^-23^. **e.** Metrics cited across all articles; investigation time refers to interaction time, sniffing time, and contact time were number of interactions refers to a unit of interaction bouts rather than a cumulative time metric; time in region of interest (ROI) includes time spent near or in a chamber or other target area. Time in ROI vs investigation time two

### Few studies investigate long-term social memories

In addition to quantifying the familiarity levels between subject and social stimulus, we quantified the length of the social memory at the time of testing. Some studies have demonstrated that the neural mechanisms of short vs long term social memory are distinct^51,52^. In addition, there is a vast literature demonstrating that for non-social memories the neural mechanisms and circuits underlying short- and long-term memories can differ^53–55^. Therefore, we asked whether social memory studies represented these equally. We found that most of the studies tested very short-term memories (see Table 1 for definitions). The intertrial intervals used (ITI; the time between learning, encoding, or familiarization with a social stimulus, and the memory test trial) were short. Short ITI’s and low levels of familiarity result in those experiments testing short-term memories. Less than 2% of articles tested social memories that had more than 24 hours since encoding began, and almost half of the articles tested social memories shorter than an hour (**Figure 6d**). Thus, for most neuroscience research probing social memory, only short-term memories of short-term familiarity have been investigated. In addition to the implications of this gap for our fundamental understanding of the brain, filling this gap is important for understanding how to treat brain diseases that affect long- and short-term memories differently, such as Alzheimer’s and other dementias^56,57^.

### Social memory metrics focus on single variables

Investigation time generally decreases upon repeated exposures to a social stimulus and increases upon exposure to a novel social stimulus. We found that this change in investigation time is homogenously used as the readout of social memory. Of the 672 articles reviewed, only 2% quantified behaviors beyond time or frequency of investigation of the social stimulus (i.e., sniffing time, time spent in proximity, number of interactions etc.) as the behavioral metric to measure social memory (**Figure 6e**). Two studies had specific additional interaction qualifications such as latency to approach or time spent following^58,59^. Quantifying and comparing the ways in which subjects interact with social stimuli could provide insight that total investigation time may not. For example, Netser et al. quantified a variety of behavioral metrics during a social discrimination paradigm using rats. Between novel and familiar social stimuli investigation, the most significant difference was found when analyzing long bouts of investigation (>19s) as opposed to the insignificant difference amongst short bouts of investigation (<6s). Four articles compared ultrasonic vocalizations across social stimulus exposures^60–63^, one quantified urine markings in response to a novel vs familiar stimulus^64^, and one measured the rate of pregnancy termination in the presence of an unfamiliar vs. a familiar mate^48^. Kavaliers measured the effects of familiarity with social odors on risk taking behavior and mate preference^65,66^. Finally, Cordero and Sandi used maintenance of social hierarchies through competition outcomes as a measure of social recognition.

### Social recognition between two familiar conspecifics and social identity are understudied

The limitation in assays and behavioral metrics have led to the social memory field almost exclusively studying social memory via discrimination of a single familiar animal from a novel animal. We found only one study that measured social memory mechanisms during the discrimination of two familiar animals. This article tested the discrimination between two familiar stimuli by different behavioral outputs from the subject; van Wingerden and van den Bos used rate of risk-taking behavior to distinguish between a “trustworthy” vs “untrustworthy” stimulus (defined as conspecifics associated with different reward probabilities^26^). Since we performed our meta-analysis, a new study was published showcasing a novel go/ no-go task where subject mice discriminate between two familiar mice to obtain a reward^25^. This design allowed the authors to study the neural dynamics of social identity encoding. We foresee that creative assays, such as these, will propel the field forward in our mechanistic understanding of social memory by dissociating mechanisms of novelty preferences from social identity encoding and recalling social memories of familiar individuals.

## Discussion

Our results indicate that social memories studied in the past two decades are biased towards short-term memories of short-term familiarity. This bias has led to underrepresentation of long-term relationships that are relevant across animal models such as kin, cagemates, members of their hierarchies and mates. Our results also show that there is little research on the neural mechanisms of long-term social memories. This is of particular importance, given that neuropathologies that affect memories, such as Alzheimer’s, tend to disrupt short-term and long-term memory in different stages of the disease^56,57^. Understanding the circuits underlying short vs long-term social memory may design and inform therapies for distinct deficits. We recognize that the exact design of a study and which social factors are included as a variable will depend on the goal of the scientific study and more behavioral testing is not always better. The goal of the study must be considered to choose the experimental design, but awareness of gaps, biases and factors that affect social behavior will help optimize future experiments in social memory. Below we detail potential directions and metrics to increase our understanding of social memories and what information we can gain in future studies.

### Incorporating long-term familiarity memories

Our results show that most social memory studies focus on short-term familiarity, with few studies involving long-term social memory. Using classical social recognition assays, limited exposure to social stimuli (15 min or less), results in social recognition that can last up to an hour in individually housed rodents^19,67^ and up to 7 days for group-housed rodents^33^. These studies suggest that it may take long-term familiarity to facilitate the long-term encoding of social identity. These differences in how long the social memories last based on exposure time, suggest the possibility of differences in mechanisms for social memories of short- vs long-term familiarity. A large body of literature shows mechanistic differences between short- and long-term non-social memories^54,68,69^. There are also some reported differences in the neural mechanisms of short and long-term social memories^52,70^. Importantly, a social memory with short-term familiarity (e.g. 10 min) could be studied as a short or a long-term memory by simply varying the interval between the acquisition vs recall. On the other hand, long-term familiarity social memories, such as memories of cagemate identity, are long-term memories that get updated as new interactions occur. For improving our fundamental knowledge of the mechanisms of social memory, the field would benefit from future studies comparing the neural mechanisms of short vs long-term social memories, as well as the comparison of short vs long-term familiarity. Incorporating the use of social stimuli with long-term ethologically-relevant relationships can facilitate the study of long-term social memory. In addition, to combat the challenges of short-term familiarity memories being forgotten, operant-based assays^17,25,26^ could make social memories of individuals with short-term familiarity more salient by pairing them with rewarding or aversive outcomes.

### Operant based social memory assays to study social identity

Our results show that there is little work to investigate how social identity memories may be represented. Relying only on investigation time, the most common behavior metric used, limits research to novelty discrimination (e.g. discrimination between a novel vs familiar animal). If social recognition is not detected with a novelty preference-based assay, is there a failure to discern and/or prefer the novelty of an animal or a failure to recognize the familiarity of a familiar one? With classical novelty-based social memory assays it is not possible to distinguish between these two possibilities. Assays that do not rely on novelty preferences allow dissociating between individual recognition and novelty discrimination. In particular, operant-based assays allow studying behavioral discrimination between multiple familiar individuals and social identity encoding (see Table 3 and **Figure 1**).

**Table 3.** Diverse social memory metrics and their potential benefits for future studies.

| Additional metrics for social memory | Potential benefit in social memory studies |
| --- | --- |
| Long vs short bouts of social investigation | Long bouts of investigation are more common with novel social stimuli, compared to short bouts (Netser et al., 2020). This distinction could improve signal to noise in social recognition measurements.<br>This metric could help in rodent models of neuropsychiatric social deficits to identify interventions that could improve social recognition. |
| Operant based tasks | Allows studying social identity and the social discrimination between two familiar individuals (Gheusi et al., 1994; Kong et al., 2023; van Wingerden & van den Bos, 2015). This is currently not possible to do with assays that rely on discrimination of familiar and novel animals.<br>May facilitate the study of long-term social memories as pairing rewards or aversive stimuli with social stimuli increases salience of the social stimuli.<br>Operant based metrics could help in our fundamental understanding of social identity and how its mechanisms differ from novelty preference. In addition, these tasks can be helpful to model disrupted discrimination among multiple familiar individuals in diseases that affect memory, such as Alzheimer's and other Dementias. |
| Combining social investigation of familiar and novel social stimuli with neurophysiological recordings | Allows potentially decoding social identity or novelty vs familiarity from neural activity measurements. This approach was recently used to describe how the hippocampal CA2 encodes social identity differently in novel vs familiar individuals<sup>89</sup>. Overall, this approach may facilitate identifying brain regions involved in social identity encoding and long-term social memory to improve our fundamental knowledge of social memory. |
| Heat-map based analysis of investigation of multiple familiar stimuli vs novel agent in SocioBox | The SocioBox assay requires subjects to discriminate a novel stimulus from 4 familiar mice which increases the social memory load on the assay, as compared to other traditional assays. Social recognition is measured using a heat map-based comparison of location and creating data-driven regions of interest which improves the signal to noise in social recognition indices<sup>27</sup>. Therefore, this assay may be more sensitive in identifying disease-related changes when studying rodent models for diseases. |
| Ultrasonic vocalizations (USV) | USV rate decreased in female mice when interacting with familiar same-sex individuals (D'Amato & Moles, 2001; Gracceva et al., 2009; Moles et al., 2007). Measure USV can be a good alternative metric in female rodents considering they have lower baseline levels of social investigation.<br>Rodent models with genetic disruptions linked to autism spectrum disorder (ASD) show decrease USVs<sup>90,91</sup>. Future studies focused on ASD rodent models could measure social memory associated USVs for additional metrics to study multiple social memory related variables in search of potential treatments and disease related mechanisms. |

### Additional metrics of social memory

A recent review discusses the limitations of the three-chamber social assays and a potential integrative, multimodal approach with simultaneous behavioral and physiological measurements at high temporal resolution^21^. In addition, recent studies have shown that short vs long bouts of social investigation are of relevance for social recognition for both mice and rats^23,71^. Both species have longer bouts of investigation with the novel stimuli, providing an additional metric beyond total investigation time. Our systematic review identified four studies that used ultrasonic vocalizations (USV) as a metric for social memory in female mice of different strains^60–63^. In these studies, USV rate decreased when interacting with familiar same-sex individuals. Another study identified in our systematic review used a data-driven approach to determining regions of interest in a social recognition test, and using this approach compared to the traditional user-defined region of interest improved the signal to noise in social recognition indices^27^. Using diverse social memory metrics could be important for identifying potential treatments for social deficits in mouse models for disease. It is possible that only some metrics may be differentially affected by disease states and thus those metrics could serve for studying potential rescue mechanisms. On the other hand, interventions that affect multiple metrics of social memory, rather than just one, could have more potential for translation. We summarize potential benefits of using additional behavioral metrics in Table 3.

### Considering biological factors in subject and social stimuli that may affect social memory

Our data also demonstrates a chronic lack of reporting of information regarding social stimuli utilized, which poses challenges for reproducibility. The omission of these details prevents the understanding of any potential differences in behavior elicited by the specifics of social stimulus identity such as sex, strain, and housing. Recent studies have shown that social stimuli state and behavior affect subject behavior. Active behavior by the social stimulus is necessary for social recognition^38^, stimuli movement can either enhance or deter social investigation from the subject^23^, and even drug administration to a constrained stimuli can effect sociability of a freely moving subject^72^, all of which highlight the importance of the two-way interaction. These studies suggest that the behavioral state (isolation, stress, age, etc.) of the social stimulus could affect the behavior of the subject and potentially the social memory metrics.

Other important factors that can impact the ethological and biological relevance of social relationships are housing condition, sex, and age. Our data demonstrate that these factors are often not reported or considered, yet they can be crucial for interpretation of results as they affect social interactions in rodents (see Table 2). Our results also show that adult male subjects dominate the social memory literature. A recent study showed that the neural mechanisms of social recognition differ between adults and adolescent rats^73^, demonstrating the importance of studying social memory at different developmental stages. Although there is ample evidence that aging affects memory, only 2% of our surveyed articles studied social memory in aged rodents. Studying aging and how it impacts social memory mechanisms is relevant for translating basic research to treat dementia and other aging related memory problems in humans. In addition, studying both male and female animals is important for translating pre-clinical research since it has been shown that many brain disorders have different rates in women and men and symptomology can differ across sexes^74^. Lastly, we also found that very few studies considered social rank in their group housed animals. Given the reports of subordinates increased attention towards dominant animals^49^, the neural mechanisms of long-term social memory may be modulated by social rank to facilitate quick behavioral discrimination of social rank.

In summary, we have quantified the volume of social memory literature utilizing different methodologies, outlined how different social and biological factors affect social memory, and the limitations with which social memory has been measured. We hope to provide context for considerations for future research design to better understand the neural mechanisms of social memory.

## Supporting information

Supplemental Table 1: Full systematic review dataset

## Acknowledgements

This work was supported by R00MH124435 and by a Burroughs Wellcome Fund Postdoctoral Enrichment fund. Behavioral schematics in the figures were created with Biorender.

## Author contributions

N.P.C. conceived the study. N.P.C. and M.C. planned and organized the study and wrote the manuscript. M.C. performed and managed the data collection and analysis. J.S.P., E.W., N.L., R.L.I., E.S.W. and A.R.C. performed data collection. M.C., A.L. and R.L.I. wrote software for data analysis. All authors reviewed the manuscript.

## Availability of Data and Materials

All raw data utilized for this article is available in the supplementary file provided.

## References

1. Alper, T. G. & Korchin, S. J. Memory for socially relevant material. J. Abnorm. Soc. Psychol. 47, 25–37 (1952).

2. Witryol, S. L. & Kaess, W. A. Sex differences in social memory tasks. J. Abnorm. Psychol. 54, 343–346 (1957).

3. Carr, W. J., Yee, L., Gable, D. & Marasco, E. Olfactory recognition of conspecifics by domestic Norway rats. J. Comp. Physiol. Psychol. 90, 821–828 (1976).

4. Porter, R. H., Wyrick, M. & Pankey, J. Sibling Recognition in Spiny Mice (Acomys cahirinus). Behav. Ecol. Sociobiol. 3, 61–68 (1978).

5. Hurst, J. L. Behavioural variation in wild house mice Mus domesticus Rutty: A quantitative assessment of female social organization. Anim. Behav. 35, 1846–1857 (1987).

6. Calhoun, J. B. Death Squared. Proc. R. Soc. Med. 66, 80–88 (1973).

7. Schweinfurth, M. K. The social life of Norway rats (Rattus norvegicus). eLife 9, e54020 (2020).

8. Barnard, C. J., Hurst, J. L. & Aldhous, P. Of mice and kin: the functional significance of kin bias in social behaviour. Biol. Rev. Camb. Philos. Soc. 66, 379–430 (1991).

9. Brown, R. Z. Social Behavior, Reproduction, and Population Changes in the House Mouse (Mus musculus L.). Ecol. Monogr. 23, 218–240 (1953).

10. Hurst, J. L. Urine marking in populations of wild house mice Mus domesticus rutty. I. Communication between males. Anim. Behav. 40, 209–222 (1990).

11. So, N., Franks, B., Lim, S. & Curley, J. P. A Social Network Approach Reveals Associations between Mouse Social Dominance and Brain Gene Expression. PLOS ONE 10, e0134509 (2015).

12. Wang, F. et al. Bidirectional control of social hierarchy by synaptic efficacy in medial prefrontal cortex. Science 334, 693–697 (2011).

13. Williamson, C. M., Franks, B. & Curley, J. P. Mouse Social Network Dynamics and Community Structure are Associated with Plasticity-Related Brain Gene Expression. Front. Behav. Neurosci. 10, (2016).

14. Beery, A. K. & Zucker, I. Sex Bias in Neuroscience and Biomedical Research. Neurosci. Biobehav. Rev. 35, 565–572 (2011).

15. Carlesimo, G. A. & Oscar-Berman, M. Memory deficits in Alzheimer’s patients: A comprehensive review. Neuropsychol. Rev. 3, 119–169 (1992).

16. Sasaguri, H. et al. APP mouse models for Alzheimer’s disease preclinical studies. EMBO J. 36, 2473–2487 (2017).

17. Gheusi, G., Bluthé, R.-M., Goodall, G. & Dantzer, R. Social and individual recognition in rodents: Methodological aspects and neurobiological bases. Behav. Processes 33, 59–87 (1994).

18. Moy, S. S. et al. Sociability and preference for social novelty in five inbred strains: an approach to assess autistic-like behavior in mice. Genes Brain Behav. 3, 287–302 (2004).

19. Ferguson, J. N. et al. Social amnesia in mice lacking the oxytocin gene. Nat. Genet. 25, 284–288 (2000).

20. Thor, D. H. & Holloway, W. R. Social memory of the male laboratory rat. J. Comp. Physiol. Psychol. 96, 1000–1006 (1982).

21. Jabarin, R., Netser, S. & Wagner, S. Beyond the three-chamber test: toward a multimodal and objective assessment of social behavior in rodents. Mol. Autism 13, 41 (2022).

22. Azahara, O., Antonio, F.-R., Felix, L. & Steven, S. A. Hippocampal CA2 sharp-wave ripples reactivate and promote social memory. Nature 587, 264–269 (2020).

23. Netser, S. et al. Distinct dynamics of social motivation drive differential social behavior in laboratory rat and mouse strains. Nat. Commun. 11, 5908 (2020).

24. Lopez-Rojas, J., de Solis, C. A., Leroy, F., Kandel, E. R. & Siegelbaum, S. A. A direct lateral entorhinal cortex to hippocampal CA2 circuit conveys social information required for social memory. Neuron 110, 1559–1572.e4 (2022).

25. Kong, E., Lee, K.-H., Do, J., Kim, P. & Lee, D. Dynamic and stable hippocampal representations of social identity and reward expectation support associative social memory in male mice. Nat. Commun. 14, 2597 (2023).

26. van Wingerden, M. & van den Bos, W. CAN YOU TRUST A RAT? USING ANIMAL MODELS TO INVESTIGATE THE NEURAL BASIS OF TRUST-LIKE BEHAVIOR. Soc. Cogn. 33, 387–413 (2015).

27. Krueger-Burg, D. et al. The sociobox: A novel paradigm to assess complex social recognition in male mice. Front. Behav. Neurosci. 10, (2016).

28. Karlsson, S. A., Haziri, K., Hansson, E., Kettunen, P. & Westberg, L. Effects of sex and gonadectomy on social investigation and social recognition in mice. BMC Neurosci. 16, (2015).

29. Markham, J. A. & Juraska, J. M. Social recognition memory: Influence of age, sex, and ovarian hormonal status. Physiol. Behav. 92, 881–888 (2007).

30. Bluthé, R. M. & Dantzer, R. Social recognition does not involve vasopressinergic neurotransmission in female rats. Brain Res. 535, 301–304 (1990).

31. Bianchi, E., Menicacci, C. & Ghelardini, C. Dual effect of morphine in long-term social memory in rat. Br. J. Pharmacol. 168, 1786–1793 (2013).

32. Gao, X., Elmer, G., Adams-Huet, B. & Tamminga, C. Social memory in mice: Disruption with an NMDA antagonist and attenuation with antipsychotic drugs. Pharmacol. Biochem. Behav. 92, 236–242 (2009).

33. Kogan, J. H., Frankland, P. W. & Silva, A. J. Long-term memory underlying hippocampus-dependent social recognition in mice. Hippocampus 10, 47–56 (2000).

34. Ehrhardt, A., Wang, B., Leung, M. J. & Schrader, J. W. Absence of M-Ras modulates social behavior in mice. BMC Neurosci. 16, 1 (2015).

35. Liu, D. et al. A Hypothalamic Circuit Underlying the Dynamic Control of Social Homeostasis. 2023.05.19.540391 Preprint at 10.1101/2023.05.19.540391 (2023).

36. Mansk, L. M. Z., Jaimes, L. F., Dias, T. L. & Pereira, G. S. Social recognition memory differences between mouse strains: On the effects of social isolation, adult neurogenesis, and environmental enrichment. Brain Res. 1819, 148535 (2023).

37. George, A., Padilla-Coreano, N. & Opendak, M. For neuroscience, social history matters. Neuropsychopharmacology 1–2 (2023) doi:10.1038/s41386-023-01566-8.

38. de la Zerda, S. H. et al. Social recognition in laboratory mice requires integration of behaviorally-induced somatosensory, auditory and olfactory cues. Psychoneuroendocrinology 143, 105859 (2022).

39. Jacobs, S., Wei, W., Wang, D. & Tsien, J. Z. Importance of the GluN2B carboxy-terminal domain for enhancement of social memories. Learn. Mem. 22, 401–410 (2015).

40. Jacobs, S. A. & Tsien, J. Z. Genetic overexpression of NR2B subunit enhances social recognition memory for different strains and species. PLoS ONE 7, (2012).

41. Tanaka, K., Osako, Y. & Yuri, K. Juvenile social experience regulates central neuropeptides relevant to emotional and social behaviors. Neuroscience 166, 1036–1042 (2010).

42. Choleris, E. et al. An estrogen-dependent four-gene micronet regulating social recognition: A study with oxytocin and estrogen receptor-α and -β knockout mice. Proc. Natl. Acad. Sci. U. S. A. 100, 6192–6197 (2003).

43. Imwalle, D. B., Scordalakes, E. M. & Rissman, E. F. Estrogen receptor alpha influences socially motivated behaviors. Horm Behav 42, 484–91 (2002).

44. Tang, A. C. et al. Effects of long-term estrogen replacement on social investigation and social memory in ovariectomized C57BL/6 mice. Horm. Behav. 47, 350–357 (2005).

45. Laham, B. J., Diethorn, E. J. & Gould, E. Newborn mice form lasting CA2-dependent memories of their mothers. Cell Rep. 34, (2021).

46. Morais, L. H. et al. Early-life oxytocin attenuates the social deficits induced by caesarean-section delivery in the mouse. Neuropsychopharmacology 46, 1958–1968 (2021).

47. Yang, E.-J., Ahn, S., Lee, K., Mahmood, U. & Kim, H.-S. Early behavioral abnormalities and perinatal alterations of PTEN/AKT pathway in valproic acid autism model mice. PLoS ONE 11, (2016).

48. Wersinger, S. R., Temple, J. L., Caldwell, H. K. & Young, W. S. Inactivation of the oxytocin and the vasopressin (Avp) 1b receptor genes, but not the Avp 1a receptor gene, differentially impairs the Bruce effect in laboratory mice (Mus musculus). ENDOCRINOLOGY 149, 116–121 (2008).

49. Dwortz, M. F., Curley, J. P., Tye, K. M. & Padilla-Coreano, N. Neural systems that facilitate the representation of social rank. Philos. Trans. R. Soc. B Biol. Sci. 377, 20200444 (2022).

50. Cordero, M. I. & Sandi, C. Stress amplifies memory for social hierarchy. Front Neurosci 1, 175–84 (2007).

51. Lin, Y.-T. et al. Conditional Deletion of Hippocampal CA2/CA3a Oxytocin Receptors Impairs the Persistence of Long-Term Social Recognition Memory in Mice. J. Neurosci. 38, 1218– 1231 (2018).

52. Sakamoto, T. & Yashima, J. Prefrontal cortex is necessary for long-term social recognition memory in mice. Behav. Brain Res. 435, 114051 (2022).

53. Do-Monte, F. H., Quiñones-Laracuente, K. & Quirk, G. J. A temporal shift in the circuits mediating retrieval of fear memory. Nature 519, 460–463 (2015).

54. Izquierdo, I., Medina, J. H., Vianna, M. R. M., Izquierdo, L. A. & Barros, D. M. Separate mechanisms for short- and long-term memory. Behav. Brain Res. 103, 1–11 (1999).

55. Lzquierdo, I. et al. Differential involvement of cortical receptor mechanisms in working, short-term and long-term memory. Behav. Pharmacol. 9, 421 (1998).

56. Hulme, C., Lee, G. & Brown, G. D. A. Short-term memory impairments in Alzheimer-type dementia: Evidence for separable impairments of articulatory rehearsal and long-term memory. Neuropsychologia 31, 161–172 (1993).

57. Miller, E. Short- and long-term memory in patients with presenile dementia (Alzheimer’s disease). Psychol. Med. 3, 221–224 (1973).

58. Oliveira, V. E. D. M., Neumann, I. D. & de Jong, T. R. Post-weaning social isolation exacerbates aggression in both sexes and affects the vasopressin and oxytocin system in a sex-specific manner. Neuropharmacology 156, (2019).

59. Phillips, M. L., Robinson, H. A. & Pozzo-Miller, L. Ventral hippocampal projections to the medial prefrontal cortex regulate social memory. eLife 8, (2019).

60. D’Amato, F. R. & Moles, A. Ultrasonic vocalizations as an index of social memory in female mice. Behav. Neurosci. 115, 834–840 (2001).

61. Gracceva, G., Venerosi, A., Santucci, D., Calamandrei, G. & Ricceri, L. Early social enrichment affects responsiveness to different social cues in female mice. Behav. Brain Res. 196, 304–309 (2009).

62. Moles, A., Costantini, F., Garbugino, L., Zanettini, C. & D’Amato, F. R. Ultrasonic vocalizations emitted during dyadic interactions in female mice: a possible index of sociability? Behav Brain Res 182, 223–30 (2007).

63. Venerosi, A., Calarnandrei, G. & Ricceri, L. A social recognition test for female mice reveals behavioral effects of developmental chlorpyrifos exposure. Neurotoxicol. Teratol. 28, 466– 471 (2006).

64. Arakawa, H., Arakawa, K., Blanchard, D. C. & Blanchard, R. J. A new test paradigm for social recognition evidenced by urinary scent marking behavior in C57BL/6J mice. Behav. Brain Res. 190, 97–104 (2008).

65. Kavaliers, M. et al. Estrogen receptors alpha and beta mediate different aspects of the facilitatory effects of female cues on male risk taking. PSYCHONEUROENDOCRINOLOGY 33, 634–642 (2008).

66. Kavaliers, M., Bishnoi, I. R., Ossenkopp, K. P. & Choleris, E. Odor-based mate choice copying in deer mice is not affected by familiarity or kinship. Anim Cogn (2021) doi:10.1007/s10071-021-01550-z.

67. Lukas, M., Bredewold, R., Landgraf, R., Neumann, I. D. & Veenema, A. H. Early life stress impairs social recognition due to a blunted response of vasopressin release within the septum of adult male rats. Psychoneuroendocrinology 36, 843–853 (2011).

68. Barros, D. M., Pereira, P., Medina, J. H. & Izquierdo, I. Modulation of working memory and of long- but not short-term memory by cholinergic mechanisms in the basolateral amygdala. Behav. Pharmacol. 13, 163 (2002).

69. Vianna, M. R. M. et al. Short- and long-term memory: differential involvement of neurotransmitter systems and signal transduction cascades. An. Acad. Bras. Ciênc. 72, 353–364 (2000).

70. Lin, Y.-T. et al. Conditional deletion of hippocampal CA2/CA3a oxytocin receptors impairs the persistence of long-term social recognition memory in mice. J. Neurosci. 38, 1218–1231 (2018).

71. Netser, S., Haskal, S., Magalnik, H. & Wagner, S. A novel system for tracking social preference dynamics in mice reveals sex- and strain-specific characteristics. Mol. Autism 8, 53 (2017).

72. Heifets, B. D. et al. Distinct neural mechanisms for the prosocial and rewarding properties of MDMA. Sci. Transl. Med. 11, eaaw6435 (2019).

73. Ferrara, N. C. et al. Isolation driven changes in Iba1-positive microglial morphology are associated with social recognition memory in adults and adolescents. Neurobiol. Learn. Mem. 192, 107626 (2022).

74. Loke, H., Harley, V. & Lee, J. Biological factors underlying sex differences in neurological disorders. Int. J. Biochem. Cell Biol. 65, 139–150 (2015).

75. Ferguson, J. N., Young, L. J. & Insel, T. R. The neuroendocrine basis of social recognition. Front. Neuroendocrinol. 23, 200–224 (2002).

76. Fukushima, H. et al. Upregulation of calcium/calmodulin-dependent protein kinase IV improves memory formation and rescues memory loss with aging. J. Neurosci. 28, 9910– 9919 (2008).

77. Guan, X. & Dluzen, D. E. Age related changes of social memory/recognition in male fischer 344 rats. Behav. Brain Res. 61, 87–90 (1994).

78. Hlinák, Z. & Krejcí, I. Social recognition in male rats: age differences and modulation by MIF-I and Alaptide. Physiol. Res. 40, 59–67 (1991).

79. Komorowska-Müller, J. A., Ravichandran, K. A., Zimmer, A. & Schürmann, B. Cannabinoid receptor 2 deletion influences social memory and synaptic architecture in the hippocampus. Sci. Rep. 11, 16828 (2021).

80. Prediger, R. D. S., Batista, L. C. & Takahashi, R. N. Caffeine reverses age-related deficits in olfactory discrimination and social recognition memory in rats: Involvement of adenosine A1 and A2A receptors. Neurobiol. Aging 26, 957–964 (2005).

81. Prediger, R. D. S., De-Mello, N. & Takahashi, R. N. Pilocarpine improves olfactory discrimination and social recognition memory deficits in 24 month-old rats. Eur. J. Pharmacol. 531, 176–182 (2006).

82. Jacobs, S. A. & Tsien, J. Z. Overexpression of the NR2A subunit in the forebrain impairs long-term social recognition and non-social olfactory memory. Genes Brain Behav. 13, 376– 384 (2014).

83. Deng, X., Gu, L., Sui, N., Guo, J. & Liang, J. Parvalbumin interneuron in the ventral hippocampus functions as a discriminator in social memory. Proc. Natl. Acad. Sci. U. S. A. 116, 16583–16592 (2019).

84. Park, G. et al. Social isolation impairs the prefrontal-nucleus accumbens circuit subserving social recognition in mice. Cell Rep. 35, (2021).

85. Thor, D. H. & Holloway, W. R. Social memory of the male laboratory rat. J. Comp. Physiol. Psychol. 96, 1000–1006 (1982).

86. Gusmão, I. D. et al. Odor-enriched environment rescues long-term social memory, but does not improve olfaction in social isolated adult mice. Behav. Brain Res. 228, 440–446 (2012).

87. Shahar-Gold, H., Gur, R. & Wagner, S. Rapid and Reversible Impairments of Short- and Long-Term Social Recognition Memory Are Caused by Acute Isolation of Adult Rats via Distinct Mechanisms. PLoS ONE 8, (2013).

88. Gabriel, P., Mastracchio, T. A., Bordner, K. & Jeffrey, R. Impact of enriched environment during adolescence on adult social behavior, hippocampal synaptic density and dopamine D2 receptor expression in rats. Physiol. Behav. 226, (2020).

89. Boyle, L. M., Posani, L., Irfan, S., Siegelbaum, S. A. & Fusi, S. Tuned geometries of hippocampal representations meet the demands of social memory. 2022.01.24.477361 Preprint at 10.1101/2022.01.24.477361 (2023).

90. Berg, E. L. et al. Developmental social communication deficits in the Shank3 rat model of phelan-mcdermid syndrome and autism spectrum disorder. Autism Res. 11, 587–601 (2018).

91. Yang, M. et al. 16p11.2 Deletion Syndrome Mice Display Sensory and Ultrasonic Vocalization Deficits During Social Interactions. Autism Res. 8, 507–521 (2015).

